# HOCl-producing Electrochemical Bandages for Treating *Pseudomonas aeruginosa*-Infected Murine Wounds

**DOI:** 10.1101/2023.09.20.558698

**Authors:** Derek Fleming, Ibrahim Bozyel, Dilara Ozdemir, Judith Alvarez Otero, Melissa J. Karau, Md Monzurul Islam Anoy, Christina Koscianski, Audrey N. Schuetz, Kerryl E. Greenwood-Quaintance, Jayawant N. Mandrekar, Haluk Beyenal, Robin Patel

**Author notes:** **Corresponding author** Robin Patel, M.D., Division of Clinical Microbiology, Mayo Clinic, 200 First Street SW, Rochester, MN 55905.

## Abstract

A novel electrochemical bandage (e-bandage) delivering low-level hypochlorous acid (HOCl) was evaluated against *Pseudomonas aeruginosa* murine wound biofilms. 5 mm skin wounds were created on the dorsum of Swiss-Webster mice and infected with 10^6^ colony forming units (CFU) of *P. aeruginosa*. Biofilms were formed over two days, after which e-bandages were placed on the wound beds and covered with Tegaderm™. Mice were administered Tegaderm-only (control), non-polarized e-bandage (no HOCl production), or polarized e-bandage (using an HOCl-producing potentiostat), with or without concurrently administered systemic amikacin. Purulence and wound areas were measured before and after treatment. After 48 hours, animals were sacrificed, and wounds were harvested for bacterial quantification. Forty-eight hours of polarized e-bandage treatment resulted in mean biofilm reductions of 1.4 log_10_ CFUs/g (9.0 vs 7.6 log_10_; p = 0.0107) *vs* non-polarized controls, and 2.2 log_10_ CFU/g (9.8 vs 7.6 log_10_; p = 0.004) *vs* Tegaderm only controls. Systemic amikacin improved CFU reduction in Tegaderm-only (p = 0.0045) and non-polarized control groups (p = 0.0312), but not in the polarized group (p = 0.3876). Compared to the Tegaderm only group, there was more purulence reduction in the polarized group (p = 0.009), but not in the non-polarized group (p = 0.064). Wound closure was not impeded or improved by either polarized or non-polarized e-bandage treatment. Concurrent amikacin did not impact wound closure or purulence. In conclusion, an HOCl-producing e-bandage reduced *P. aeruginosa* in wound biofilms with no impairment in wound healing, representing a promising antibiotic-free approach for addressing wound infections.

## Introduction

The growing threat of antibiotic resistant bacterial pathogens has created a need for alternative antimicrobial strategies. This is particularly true in the case of chronic wound infections; it has been estimated that nearly 90% of wound isolates may harbor resistance to at least one antibiotic, with nearly 30% being resistant to at least six antibiotics.^1^ Among these resistant isolates, *Pseudomonas aeruginosa*, a Gram-negative pathogen that is intrinsically resistant to a variety of antibiotics and exhibits high rates of acquired resistance,^2^ is a commonly identified chronic wound pathogen.^3, 4^

Further augmenting antibiotic resistance is the ability of *P. aeruginosa* and other pathogens to form biofilms, communities of microorganisms protected by a heterogenous matrix of polysaccharides, proteins, DNA, and other molecules, termed extracellular polymeric substance (EPS). Biofilm-associated infections can be recalcitrant to current therapeutics, and impede wound closure, leading to a perpetual state of inflammation and delaying healing.^5, 6^ In the United States, nearly 7 million patients experience chronic wounds annually,^7^ with an estimated 60% of those wounds associated with microbial biofilms.^8^ Given the recalcitrance of chronic wound infections, and the common involvement of multi drug-resistant (MDR) *P. aeruginosa*, novel antibiofilm strategies that do not contribute to further resistance are necessary.

Hypochlorous acid (HOCl) is a biocide, naturally produced by phagocytes,^9, 10^ that has activity against both bacteria and fungi.^11-13^ A challenge in its clinical use as an anti-infective has been the inability to deliver it continuously. In previous work, we developed an electrochemical platform for generating HOCl *in situ*. This platform was active against both bacterial and fungal biofilms *in vitro*.^11-14^ Here, we show for the first time that an HOCl-producing electrochemical bandage (e-bandage), controlled by a customized ‘wearable’ potentiostat, effectively treats *P. aeruginosa* biofilm infections in an *in vivo* environment. We evaluated the HOCl-producing e-bandage activity against *P. aeruginosa*-infected murine wounds by quantifying viable bacterial cell reduction in the wound bed, evaluating wound healing factors (wound area reduction, purulence score, histopathology profile, and inflammatory cytokines), and measuring HOCl concentration in the wound bed. Lastly, we compared HOCl producing e-bandage treatment efficiency with amikacin treatment.

## Methods and Materials

### Electrochemical bandage

The e-bandage and wearable potentiostat are described in previous studies.^15, 16^ Briefly, the e-bandage comprises three integrated electrodes: functional circular carbon fabric working and counter electrodes, each with surface areas of 1.77 cm^2^ (Panex 30 PW-06, Zoltek Companies Inc., St. Louis, MO), along with a silver/silver chloride (Ag/AgCl) wire that functions as a quasi-reference electrode (QRE). The operational potential of the working electrode is maintained at +1.5 V_Ag/AgCl_ using a wearable potentiostat. Working and counter electrodes are separated by two layers of cotton fabric, with an additional layer placed above the counter electrode to facilitate moisture retention. To secure the fabrics, silicone adhesive is employed, which partially covers the external border of the electrodes and cotton layers. The QRE is affixed between the two cotton fabric layers that isolate the carbon electrodes. Titanium wires (TEMCo, Amazon.com, catalog #RW0524) are then connected to opposing ends of the e-bandage using nylon sew-on caps (Dritz, Spartanburg, SC, item#85). When the bandage is polarized at physiological conditions, HOCl generation occurs via the following reactions:

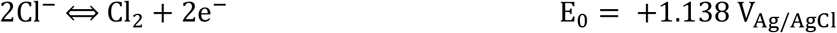

Cl_2_ (g) + H_2_O (l) = Cl^-^ (aq) + Cl_2_ (g) + H_2_O (l) = Cl^-^ (aq) + HOCl (aq) + H^+^ (aq) (pH =7.4). At pH 7.4 and 25°C, HOCl dissociates to ∼57% HOCl and ∼43% ClO^-^.

HOCl (aq) + H^+^ (aq) (pH =7.4). At pH 7.4 and 25°C, HOCl dissociates to ∼57% HOCl and ∼43% ClO^-^.

### Mice skin wound infection model

All animal experiments were approved by the Mayo Clinic Institutional Animal Care and Use Committee under protocol approval number, A00003272-20. To generate full-thickness skin wounds on Swiss Webster mice (Charles River, Wilmington, MA), animals were anesthetized by intraperitoneal injection of ketamine (90 mg/kg) and xylazine (10 mg/kg) Buprenorphine ER-Lab (1 mg/kg) administered subcutaneously for analgesia. Mature wound biofilms were generated as in our previous studies.^17, 18^ The dorsal surface was shaved and disinfected, and a 5-mm biopsy punch (Acuderm Inc., Fort Lauderdale, FL) utilized to create a circular, full thickness skin wound. Wounds were infected with 10^6^ colony forming units (CFUs) of a clinical multidrug-resistant *P. aeruginosa* IDRL-11442 isolate suspended in 0.9% sterile saline. *P. aeruginosa* IDRL-11442 is a wound isolate that exhibits ‘Difficult-to-Treat Resistance,’ in that it is resistant to first-line antibiotics piperacillin/tazobactam, cefepime, ceftazidime, meropenem, aztreonam, ciprofloxacin, and levofloxacin (although it is colistin-susceptible).^19^ Uninfected control mice were administered 10 μl of saline as a vehicle control. Bacterial suspensions were allowed to settle into wound beds for 5 minutes, after which wounds were covered with semi-occlusive transparent Tegaderm® (3M, St. Paul, MN) using liquid adhesive Mastisol® (Eloquest Health care, Ferndale, MI). Wound photographs were taken, and wound diameters recorded every other day with a Silhouette wound imaging system (Aranz Medical Ltd, Christchurch, NZ). Purulence was scored before treatment and before sacrifice to gauge immune response to biofilm infection and treatment. The scoring system was based on our previous work^17^ using the following scaling model: 0 – no exudate in the wound-bed; 1 – slight turbid exudate at the wound site; 2 – mild amount of white exudate at the wound site; 3 – moderate amount of white exudate at the wound site; 4 – moderate amount of yellowish exudate at the wound site; 5 – large amount of turbid yellow exudate extending beyond the wound-bed.

### e-Bandage treatment

After mouse wound bed infections were established for 48 hours, mice were anaesthetized, Tegaderm removed, and wearable potentiostats affixed to the skin at the scruff of the neck. Sterile e-bandages pre-hydrated in sterile 1X phosphate buffer saline (1X PBS) and 200 μl of sterile hydrogel (1.8% [w/v] xanthan gum in 1X PBS) were injected between fabric layers. Another 200 μl of hydrogel was added on top of the wound beds, and e-bandages sutured on top (two sutures, one on each lateral side of the bandage to ensure continuous contact with the wound bed with mouse activity) and attached to the female socket of the wearable potentiostats. An additional 200 μl of hydrogel was added on top of the e-bandages and the entire bandage and wound were covered with Tegaderm. 3V coin cell batteries (Ecr1220 Energizer, St. Louis, MO) were inserted into each potentiostat to start treatment (polarized e-bandage treatment). Treatment proceeded for 48 hours with hydrogel refreshment and battery changes every 24 hours. Potentials of the working electrodes relative to the QREs were measured at the start of treatment, before and after each battery change, and before euthanasia to ensure device operation. Control groups included mice with hydrogel and Tegaderm only and mice with non-polarized e-bandages (i.e., no potentiostat). Additionally, the experimental and control groups were tested with concurrent amikacin dosing. Previously, we determined the pharmacokinetic profile of amikacin in Swiss Webster mice to select a treatment dose of 15 mg/kg subcutaneous every 6 hours.^18^ At least 8 mice were included in each experimental and control group.

### Wound biofilm quantification

Following treatment, Tegaderm and e-bandages were removed from the wound bed and wound tissue excised with a 10 mm biopsy punch tool (Acuderm Inc., Fort Lauderdale, FL). Skin tissue was weighed, homogenized (Omni International, Kennesaw, GA) in sterile PBS, sonicated in for 5 minutes in a water bath, and vortexed for 30 seconds. 100 μl of the resulting homogenate was serially diluted (10-fold dilutions) in 0.9% saline and CFUs determined by spread-plating 100 μl of each dilution onto tryptic soy agar with 5% sheep blood. After 24 hours of incubation at 37°C, colonies were counted, and results reported as log_10_ CFU/g.

### Histopathology

A subset of animals (n=3) from each group underwent assessment of wound histopathology. Wound beds were removed via a 10 mm biopsy punch and fixed in 10% formalin. Fixed samples were stained with hematoxylin and eosin (H&E), and slides were blindly analyzed by a board-certified clinical pathologist for overall inflammation (0-none, 1-mild, 2-moderate, 3-severe), abscess formation (Y/N), ulceration (Y/N), tissue necrosis (Y/N), and neutrophilic infiltration (Y/N).

### Toxicity screen analysis and inflammatory panel screening

Following euthanasia, blood was obtained through cardiac puncture and subjected to centrifugation to separate it. The resulting serum samples were assessed using a Piccolo® Xpress™ Chemistry Analyzer at the Mayo Clinic Central Clinical Laboratory. The concentrations of including glucose, amylase, blood urea nitrogen, alkaline phosphatase, alanine aminotransferase, aspartate aminotransferase, gamma glutamyltransferase, lactate dehydrogenase, C-reactive protein, total bilirubin, creatinine, uric acid, albumin, total protein, calcium, chloride, magnesium, potassium, sodium, and total carbon dioxide levels were determined. Additionally, serum samples were subjected to analysis using a MesoScale Discovery SQ 120 to assess the levels of pro-inflammatory biomarkers, including IFN-γ, IL-10, IL-12p70, IL-2, IL-4, IL-5, IL-6, TNF-α, and KC/GRO.

### Statistical analysis

Initial comparisons among experimental groups were conducted through the Kruskal-Wallis test. Subsequent pairwise comparisons between groups were carried out using the Wilcoxon rank sum test. Non-parametric tests were selected due to limited sample sizes and absence of support for the assumption of a normally distributed dataset. All tests were double-tailed, with statistical significance considered for p-values below 0.05. Corrections for False Discovery Rate were performed for all comparisons with group sizes greater than three. The analytical process utilized SAS software (version 9.4, SAS Institute), with GraphPad Prism (software version 8.0, GraphPad Software) employed for generating graphs.

## Results

### Polarized e-bandages produced HOCl *in situ*

Previously, we used microelectrodes to show that these e-bandages produce HOCl at the working electrode, and that the HOCl can penetrate into biofilms and explant tissue.^15, 20^ Here, free chlorine spectrophotometer test kits (TNT866; Hach Company, Ames, IA) were used to measure total HOCl content in wounds. Wounds harvested from mice treated with polarized e-bandages had higher levels of HOCl than those treated with non-polarized e-bandages (p = 0.170) or Tegaderm alone (p = 0.0045) (**Table 1**).

**Table 1.** Endpoint HOCl content (μM) of wounds.

| Group | N | Mean | Standard deviation |
| --- | --- | --- | --- |
| Tegaderm-only | 8 | 66.5 | 16.8 |
| Polarized e-bandage | 9 | 161.7 | 59.8 |
| Non-polarized e-bandage | 10 | 80.7 | 43.2 |

### Polarized e-bandage treatment reduced bacterial load

To test if HOCl-producing e-bandage treatment reduces wound biofilm bacterial burden *in vivo*, 5-mm mouse dermal wounds were infected with *P. aeruginosa* and biofilms were allowed to establish for 48 hours before treatment. Wounds were treated with either polarized or non-polarized e-bandages and compared to Tegaderm alone. After 48 hours of treatment, endpoint wound CFUs were quantified. Treatment with polarized e-bandages resulted in lower bacterial loads than non-polarized e-bandages (p = 0.0048) or Tegaderm alone (p = 0.004, **Figure 1**).

**Figure 1.**
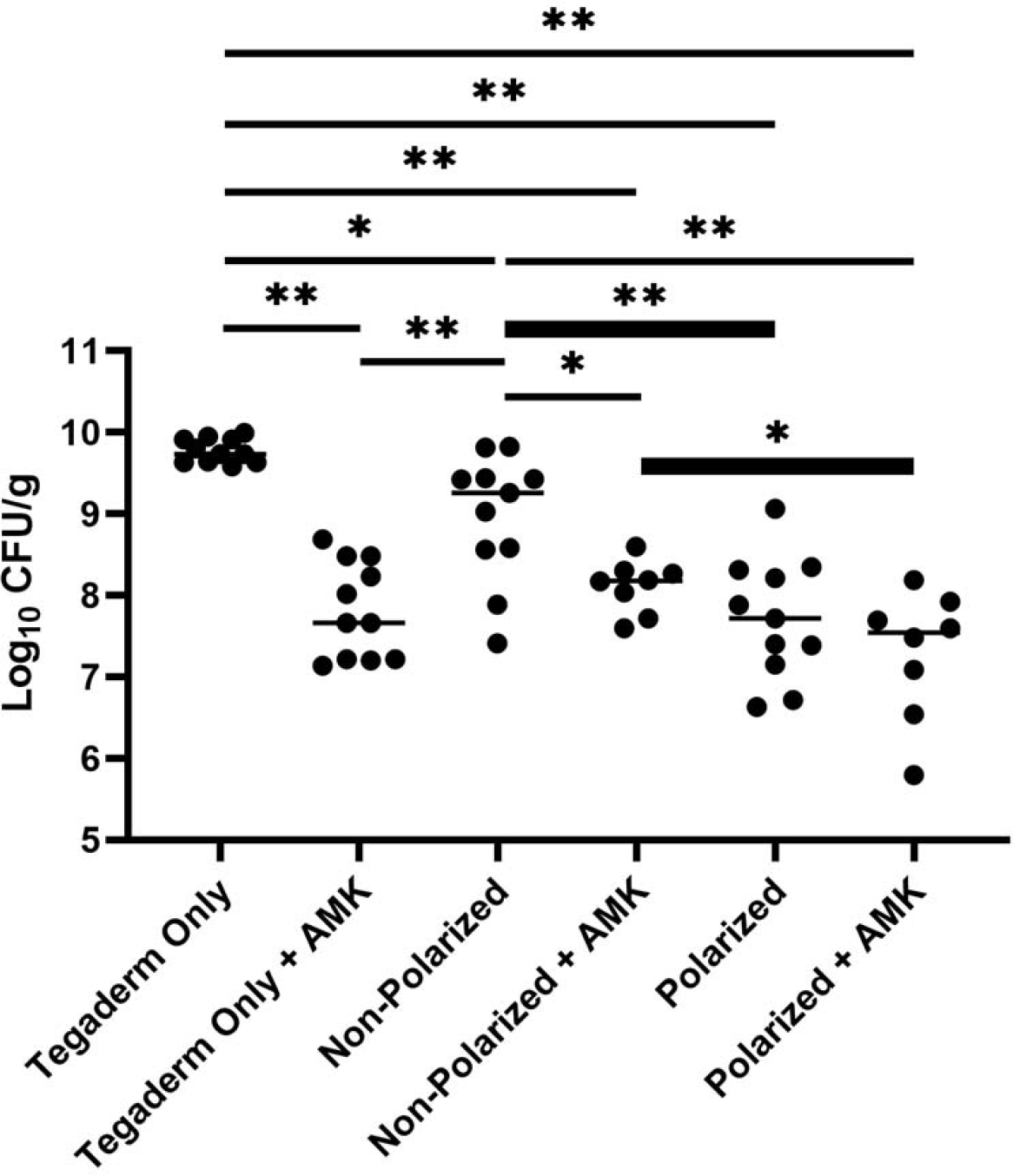
Colony forming units (CFU) counts of wounds treated with e-bandages and/or amikacin. 48-hour *P. aeruginosa* wound bed biofilms were treated for 48 hours with either polarized (HOCl-producing) or non-polarized e-bandages, with or without systemic amikacin (AMK) and compared to Tegaderm only controls, with or without systemic amikacin. Statistical analysis was performed using the Wilcoxon rank sum test with correction for false discovery rate. N ≥ 8. *p ≤0.05, **p ≤0.01

To test if the e-bandages synergize with systemic antibiotics against established wound biofilms, additional mice from all three groups were administered concurrent systemic amikacin for the duration of the e-bandage treatment window. Although amikacin alone was effective in reducing CFU counts compared to Tegaderm-only controls without amikacin (p = 0.0045), no additional reduction in CFUs was found with polarized e-bandage treatment (p = 0.312, **Figure 1**).

### Treatment of infected wounds with e-bandages did not affect wound healing

To determine the effect of e-bandage and/or amikacin treatment on wound healing, wound areas were measured before and after treatment. No significant differences in overall wound closure percentage were observed between any group (**Figure 2**).

**Figure 2.**
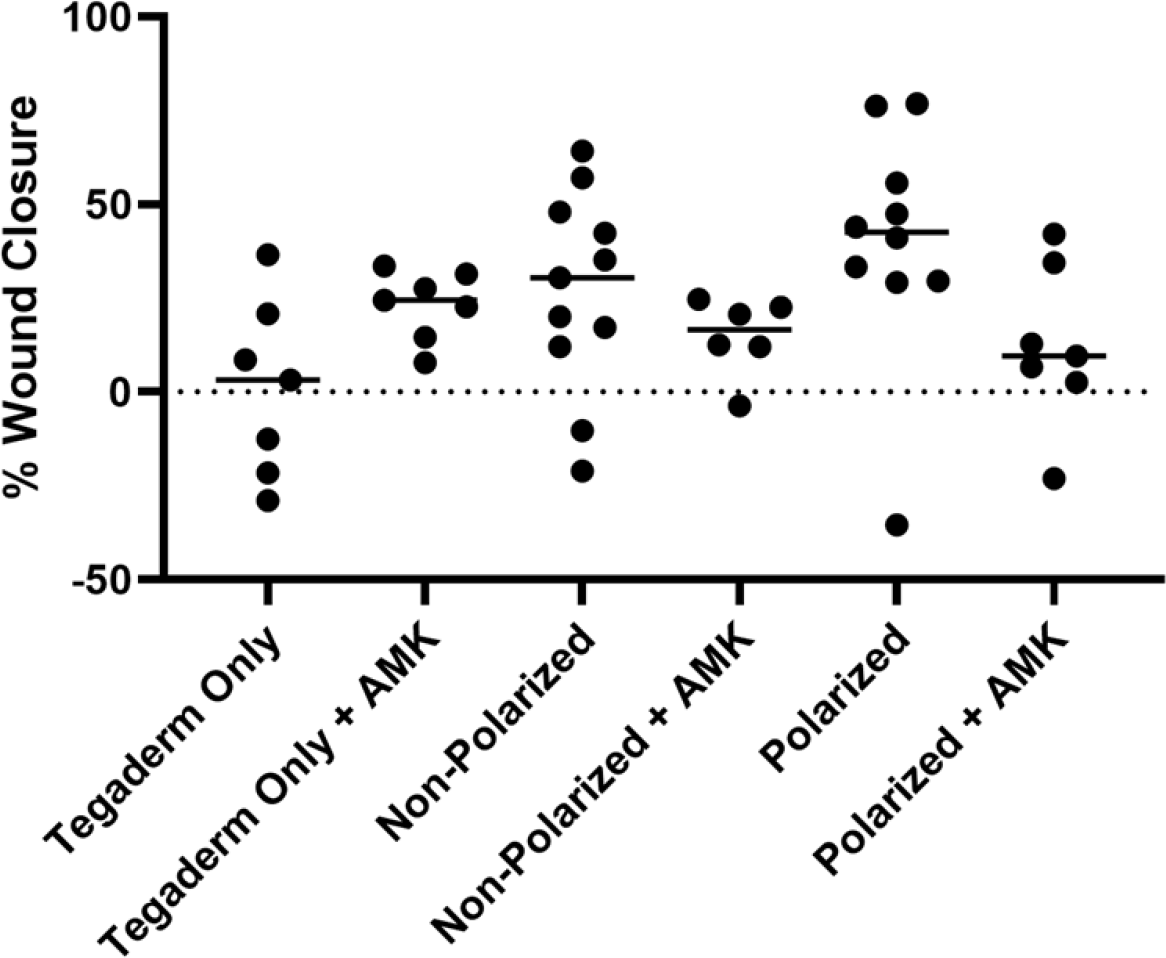
Wound closure of wounds treated with e-bandages and/or amikacin. 48-hour *P. aeruginosa* wound bed biofilms were treated with either polarized (HOCl-producing) or non-polarized e-bandages, with or without systemic amikacin (AMK) and compared to Tegaderm only controls, with or without systemic amikacin. Wound area was measured before and after 48 hours of treatment. Statistical analysis was performed using the Wilcoxon rank sum test with correction for false discovery rate. N ≥ 8.

### Treatment of infected wounds with polarized e-bandages resulted in reduced purulence

To determine the effect of e-bandage and/or amikacin treatment on purulence within the wound beds, wounds were scored for purulence before and after treatment. Treatment with polarized e-bandages led to significant improvement in purulence compared to either Tegaderm-only or non-polarized controls (p = 0.009 and 0.048, respectively). Purulence reduction within the non-polarized group was not different than in the Tegaderm-only group (p = 0.064). Concurrent amikacin did not improve purulence reduction in any group (**Figure 3**).

**Figure 3.**
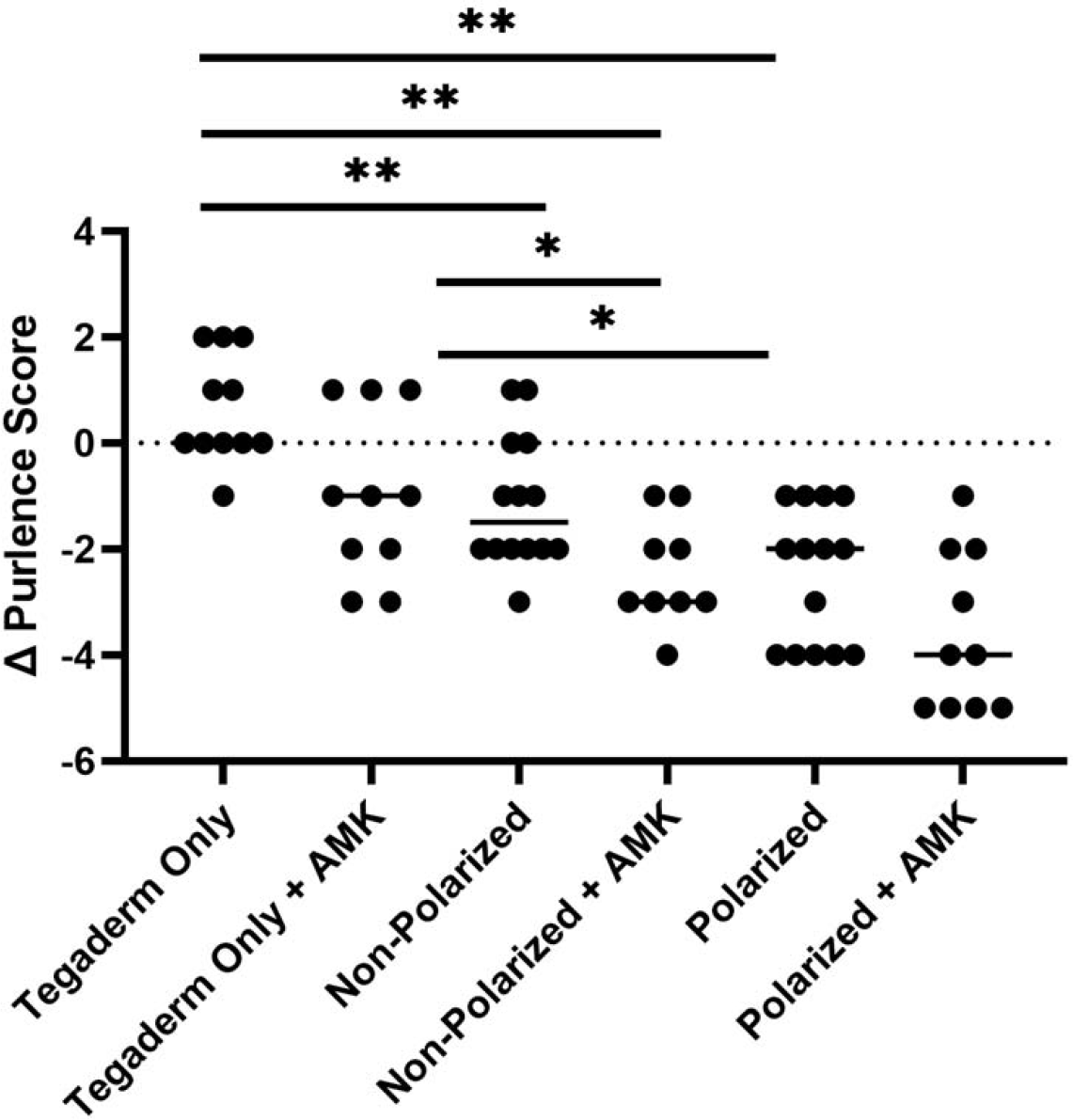
Reduction in purulence for wounds treated with e-bandages and/or amikacin. 48-hour *P. aeruginosa* wound bed biofilms were treated with either polarized (HOCl-producing) or non-polarized e-bandages, with or without systemic amikacin (AMK) and compared to Tegaderm only controls, with or without systemic amikacin. Wound purulence was scored before and after 48 hours of treatment. Statistical analysis was performed using the Wilcoxon rank sum test with correction for false discovery rate. N ≥ 8. *p ≤0.05, **p ≤0.01

### No tissue toxicity from the polarized e-bandage treatment was observed

To determine if e-bandage treatment resulted in additional tissue toxicity beyond that occurring from infection alone, three wounds from each non-antibiotic treated group were harvested and fixed in 10% formalin before being processed for H&E staining, after which a board-certified clinical pathologist blindly evaluated them (**Table 2)**. Moderate to acute inflammation of epidermal and dermal layers and ulceration were observed in all tissue samples. Abscess formation was noted in all samples except for one from the non-polarized group. The same sample was also the only one to not exhibit significant neutrophilic inflammation.

**Table 2.**
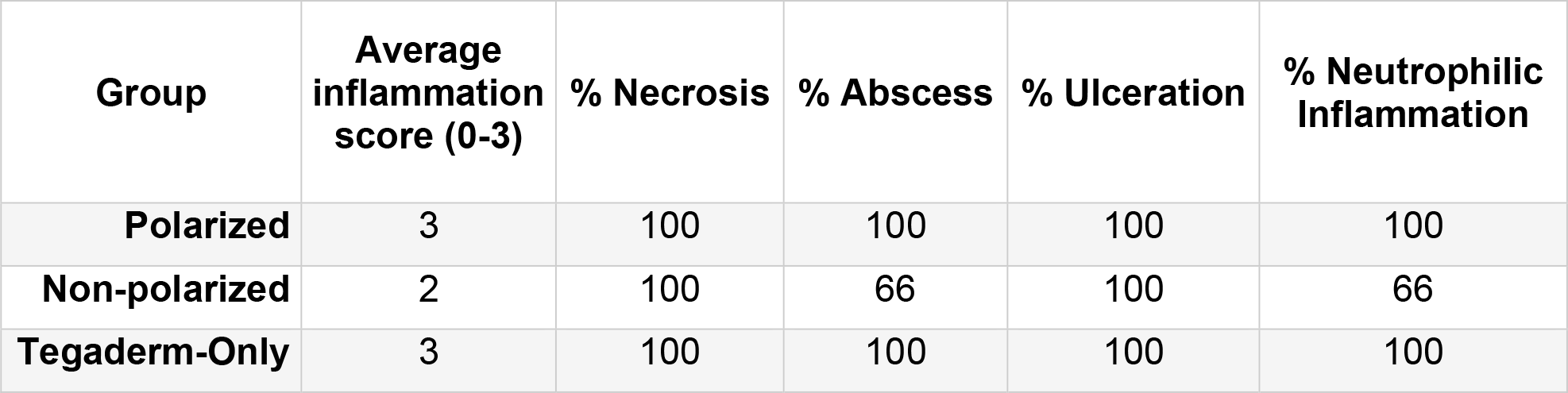
Histopathological profiles.

### Assessment of blood biomarkers

A subset (n=3) of animals treated with polarized e-bandages in comparison to the non-polarized group and Tegaderm-only controls was utilized to examine the immune response and general health of infected animals based on assessment of blood parameters. Serum was collected at the time of euthanasia and used to measure the levels of inflammatory cytokines and blood biochemistry analytes (**Table 3**). Compared to Tegaderm-only controls, the polarized group displayed subtle but statistically significant differences in serum keratinocyte chemoattractant/human growth-regulated oncogene (KC/GRO; 565.6 ±168 pg/mL vs 259.7 ±47.1 pg/mL), creatinine (0.1 ±0.1 mg/dL vs 0.4 ±0.1 mg/dL), and potassium levels (9.9 ±0.4 mmol/L vs 8.5 ±0.6 mmol/L). Similarly, the non-polarized group exhibited slight variances in serum KC/GRO (520.9 ±98.7 pg/mL vs 259.7 ±47.1 pg/mL; p = 0.0495), uric acid (1.6 ±0.5 mg/dL vs 1.0 ±0.0 mg/dL), calcium (10.7 ±0.1 mg/dL vs 10.4 ±0.2 mg/dL), and potassium levels (9.5 ±0.3 mmol/L vs 8.5 ±0.6 mmol/L) compared to the Tegaderm-only control group. There were no significant differences across all analytes between the polarized and non-polarized groups.

**Table 3.** Inflammatory cytokines and blood biochemistry analytes with significant differences between groups.

| Groups Compared |  |  | P-values |  |  |  |  |
| --- | --- | --- | --- | --- | --- | --- | --- |
| Group |  | Group | Cr | UA | Ca | K | KC/GRO |
| Tegaderm Only | vs | Non-Polarized | 0.2386 | 0.0369 | 0.0463 | 0.0463 | 0.0495 |
| Tegaderm Only | vs | Polarized | 0.0463 | 0.2207 | 0.3687 | 0.0463 | 0.0495 |
| Non-Polarized | vs | Polarized | 0.4867 | 0.0833 | 0.5127 | 0.2752 | 0.8273 |

Cr - Creatinine; UA - Uric acid; Ca - Calcium; K – Potassium; KC/GRO - Keratinocyte chemoattractant/human growth-regulated oncogene. Statistical analysis was performed using the Wilcoxon rank sum test.

## Discussion

*P. aeruginosa* is a commonly isolated chronic wound pathogen that is notorious for its intrinsic resistance and rapid acquisition of resistance to multiple classes of antibiotics.^21^ Additionally, *P. aeruginosa* is a prominent biofilm producer, further promoting its recalcitrance to traditional antibiotic treatment and clearance by the host immune system, delaying healing when present in wounds.^5, 22^ Here, a novel e-bandage was tested for the *in situ* delivery of a potent biocide, HOCl. Utilizing HOCl as a biocidal agent circumvents the issue of resistance development frequently observed with traditional antimicrobial agents. As stated in a recent publication, “There has not been a single verified claim of clinical resistance [to HOCl] over more than 100 years of careful evaluation”;^23^ this statement is corroborated by our efforts to select HOCl resistance in bacterial pathogens.^24^ Additionally, HOCl produced by the e-bandage is delivered at concentrations below the cytotoxic limit <286μM.^10^ HOCl-producing e-bandages were tested for 48 hours against 2-day *P. aeruginosa* biofilms in an *in vivo* murine dermal wound model. In confirmation of previous tests of this HOCl-producing electrochemical approach,^12, 13, 15, 24^ polarization of e-bandages resulted in more HOCl within wounds in comparison to non-polarized and Tegaderm-only controls (161 vs. 67 and 81 μM, respectively) after 48 hours of treatment.

HOCl-producing e-bandages reduced *P. aeruginosa* populations within wound beds. 48 hours of polarized treatment resulted in average bacterial loads 2.2 log_10_ CFU/g lower than Tegaderm-only controls (p = 0.004), and 1.4 log_10_ CFU/g lower than non-polarized bandages (p = 0.0048). Combinatorial treatment with e-bandages and amikacin was also tested, with no added microbicidal activity when compared to amikacin alone or polarization alone. Combinatorial treatment with amikacin and polarized e-bandages did, however, result in reduced bacterial counts compared to non-polarized e-bandages with amikacin (p = 0.0195). Considering that Tegaderm-only controls harbored more CFUs than non-polarized controls, likely due to physical disturbance of the biofilm and biofilm drying due to increased air volume underneath the Tegaderm layer, the greater microbicidal activity observed in polarized vs non-polarized amikacin-treated mice could indicate augmentation of antibiotic activity.

Unlike our previous work investigating hydrogen-peroxide- (H_2_O_2_-) producing e-bandages,^18^ HOCl-producing e-bandages did not accelerate wound healing over 48 hours of application, indicating that H_2_O_2_ might be more effective at augmenting wound healing. Devices that produce both HOCl and H_2_O_2_ could be explored in the future. It should be noted that HOCl e-bandage treatment did result in a significant reduction in wound purulence compared to both Tegaderm-only and non-polarized controls (p = 0.009 and 0.048, respectively), similar to findings with H_2_O_2_-producing e-bandages.^18^ This indicates that the overall severity of infection after e-bandage treatment has been reduced, resulting in a tempered inflammatory response and more favorable conditions for wound healing. One possible explanation is that 48 hours of treatment was insufficient to observe improvement in wound closure by these devices.

As expected, concentrations of HOCl produced by the e-bandages showed no apparent toxicity within the mouse model. Blind review of wound sections from polarized, non-polarized, and Tegaderm-only groups revealed no significant differences in overall inflammation, abscess formation, ulceration, tissue necrosis, or neutrophilic infiltration between infected tissues in the different groups. Furthermore, an assay of inflammatory cytokines and blood biochemical analysis revealed that, in comparison to Tegaderm-only controls, there were minor variations in serum KC/GRO, creatinine, and potassium levels with polarized e-bandage treatment. The non-polarized group also exhibited slight variances in serum KC/GRO, uric acid, calcium, and potassium levels compared to the Tegaderm-only control group, suggesting that, if there is any effect, it is the bandage itself, not HOCl production, that is the cause of variation. Importantly, no significant differences were found between the polarized and non-polarized groups across all analytes. It should be noted that throughout the course of the experiments, minor to moderate discoloration of the skin surrounding the wound was observed with polarized treatment. Discolored matter was physically removable by cleansing with an alcohol swab, suggesting that it related to bandage debris.

While the findings outlined in this study show promise, there are several limitations. Only a single treatment timepoint and a single bacterial strain was evaluated. Extension of duration of treatment should be explored in the future; this could hypothetically improve antibacterial efficacy, as well as overall wound healing. Furthermore, future endeavors could be aimed at enhancing the overall performance of the e-bandage system by integrating hydrogels that enhance moisture retention, essential for optimal e-bandage operation, as well as by assessing alternative electrolytes that may augment HOCl production at comparable potentials. Concurrent treatment with other systemic and/or topical antimicrobial agents could be explored. Finally, sharp debridement of the wound prior to treatment might improve e-bandage efficacy, as mature, densely populated biofilms are generally more difficult to eradicate than thinner, less populated biofilms, and treatment of thinner biofilms may allow for greater delivery of HOCl to the entirety of the biofilm matrix.

In conclusion, HOCl-producing e-bandages effectively reduced *P. aeruginosa* biofilms in a murine dermal wound infection model, with minimal evidence of toxicity and reduced purulence. Based on these results, HOCl-producing e-bandages are a promising strategy to treat *P. aeruginosa* wound infections and circumvent multi-drug resistance.

## Acknowledgments

Research reported in this publication was supported by the National Institute of Allergy and Infectious Diseases of the National Institutes of Health under award number R01AI091594.

